# Topological impact of negative links on the stability of resting-state brain network

**DOI:** 10.1101/2021.01.07.425720

**Authors:** Majid Saberi, Reza Khosrowabadi, Ali Khatibi, Bratislav Misic, Gholamreza Jafari

**Author notes:** **Corresponding author:** Reza Khosrowabadi, **Address:** Institute for Cognitive and Brain Science, Shahid Beheshti University, Evin Sq., Tehran19839-63113, Iran, **Email**.

## Abstract

Stability is a physical attribute that stands opposite the change. However, it is still unclear how the arrangement of links called topology affects network stability. In this study, we tackled this issue in the resting-state brain network using structural balance. Structural balance theory employs the quality of triadic associations between signed links to determine the network stability. In this study, we showed that negative links of the resting-state network make hubs to reduce balance-energy and push the network into a more stable state compared to null-networks with trivial topologies. In this regard, we created a global measure entitled ‘tendency to make hub’ to assess the hubness of the network. Besides, we revealed nodal degrees of negative links have an exponential distribution that confirms the existence of negative hubs. Our findings indicate that the arrangement of negative links plays an important role in the balance (stability) of the resting-state brain network.

## 1. Introduction

The brain has assumed as a complex network that its regions are structurally or functionally connected. The non-trivial characteristics of the complex brain network enable it to process and communicate neural information efficiently and produce cognitive functions and complex behaviors [1–4].

In recent years, brain researchers applied the state of the art of physics to study complex brain networks. For instance, Dante R. Chialvo showed that the large-scale resting-state networks are located at critical states which means they always stay close to the phase transition [5–6]. Besides, Enzo Tagliazucchi et.al. also discovered that loss of consciousness pushes the networks from the critical states to stable states [7].

In general, a system is called critical when it tends to make transitions between different states. Criticality is the opposite side of stability. According to the principle of minimum energy, a physical system loses energy and leaves the critical state toward a stable state. Also, a physical system remains stationary in a stable state with minimum energy level until receiving external energy.

To explore the stability of the functional brain networks, scientists usually divide regional brain activations into shorter temporal segments, extract functional connectivity of each segment, then explore the variability of functional connections over the segments and consider the inverse of them as the stability [8–10]. In the following, we explain two major weaknesses of this procedure:

First, as fMRI time courses have a low signal-to-noise ratio, when we divide them into shorter time segments, the validity of extracted functional connections can be questionable [8]. Moreover, the correlation coefficient as a common measurement of the functional connection is also vulnerable to low numbers of time points [11-12]. So the observed temporal variations of the functional connections may be a consequence of systematic errors.

Second, the process ignored the emergent property of complex brain networks and mainly determines the stability of each functional link separately then aggregates them to explain network stability. The emergent property describes that parts of a complex system don’t work individually and belong to a whole [13] and the whole is greater than the sum of the parts [14]. Therefore, we have to consider the links interrelatedly and simultaneously as a whole to respect the complexity of the brain network.

To overcome the above-mentioned shortages, we decided to employ the structural balance theory to assess the stability of functional brain networks. The structural balance is a well-known approach to investigate the stability of complex social networks [15–17]. It does not restrict us to segment time courses and has no conflict with the emergent property of the brain network. Actually, balance theory investigates the association between two entities in presence of a third-party, this is in agreement with emergent property.

The structural balance theory refers to Fritz Heider’s researches on personal interrelations [18]. The theory classified the quality of relationships between three entities as balanced and imbalanced (Fig. 1.a). Balance triads: “a friend’s friend is a friend” and “an enemy’s enemy is a friend”; Imbalanced triads: “a friend’s friend is an enemy” and “an enemy’s enemy is an enemy” [19]. Entities of an imbalanced triad are frustrated about their conditions and try to change their relationships to reach one of the balanced conditions. This is analogous to the transition of an unstable physical system toward a stable state with a lower energy level based on the principle of minimum energy. Accordingly, it can be assumed that a balanced (stable) triad situates stationary in a low-level energy state despite an imbalanced (unstable) triad that tries to solve its frustrations to reduces its energy and moves to the stable states [20].

**Figure 1:**
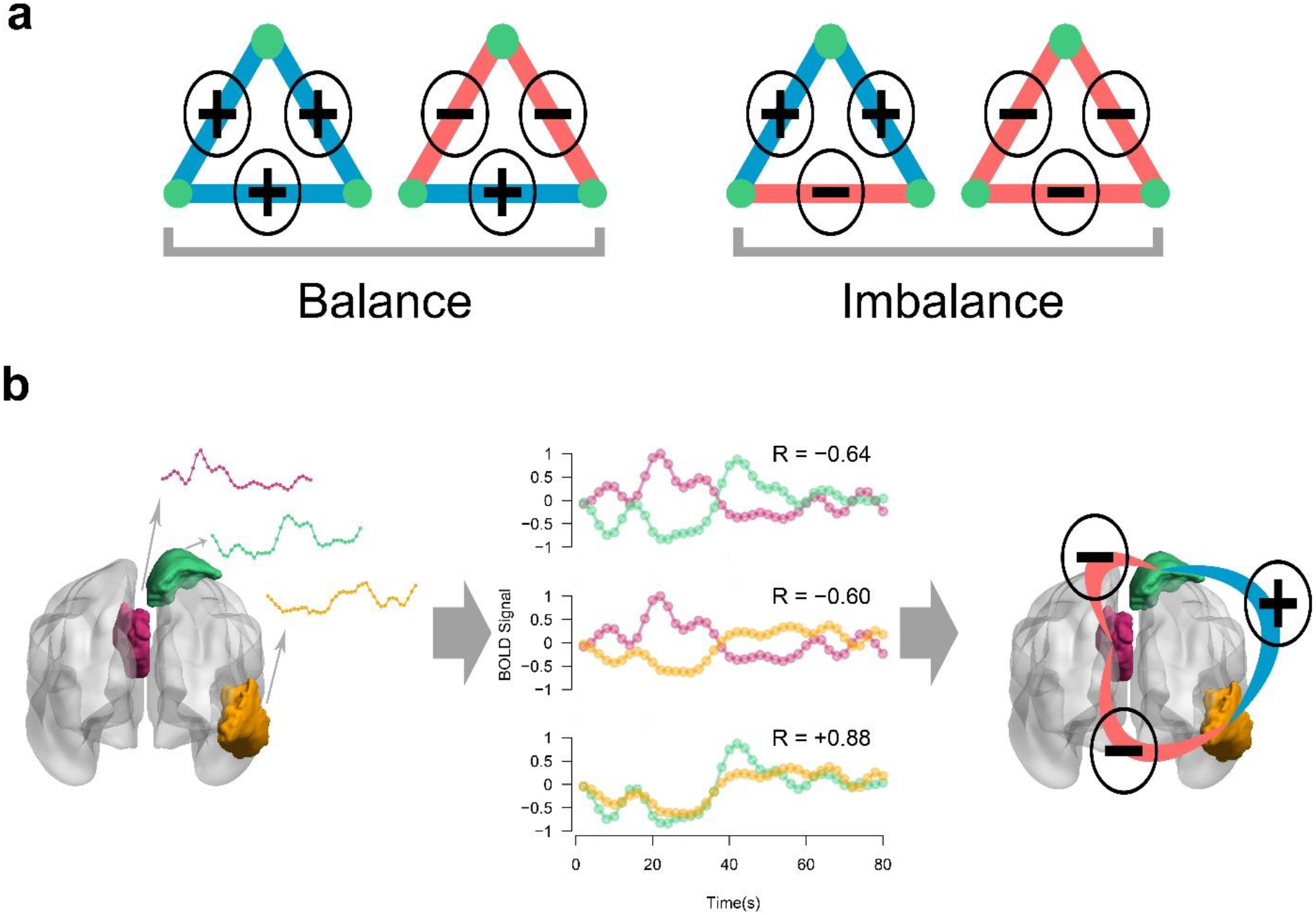
Triadic associations in balance theory. (**a**) Possible triad types of a signed network. Blue and red colors denote positive and negative links. (**b**) Formation of a triadic relation in the brain. Activity pattern of three brain regions (left) correlate together (middle) and build a balanced triangle (right). The selected regions of interest are colored violet, green, and orange that belong to Default Mode Network, Dorsal Attention Network, and Visual network of the brain, respectively. Abbreviations: R - Pearson correlation coefficient.

Also, structural balance can be investigated in a signed network where positive and negative links represent friendship and hostility, respectively [21]. Consequently, the balance of a signed network is determined by counting the number of balanced and imbalanced elements. In this way, we encounter a large number of triads, quartets, and higher-order cyclic relations and their balances [15]. Since the tension of a frustrated cycle is reduced by increasing its length, we can derive a first-order approximation of the balance by considering only the triadic relations of a signed network. In this regard, Marvel et.al. introduced the concept of energy of the signed network [22]. They formulated the balance-energy of a signed network as the difference between the number of balanced and the number of imbalanced triads. In this context, the topology of signed links represents the network state and a state is stable if and only if all of its triads are balanced. A stable network has the least energy and remains stationary since there is no dynamic demand to change link signs due to the presence of imbalanced triads.

Until here we described the importance of brain network stability and offered balance theory as a proper assessment approach for it. Now, let’s address the main question of this research. In recent years, brain scientists have devoted a lot of effort to investigate the topological aspects of brain networks and found that topological properties are the key factors in brain functions and affect behavioral and cognitive functionalities [23-26], but there is no outstanding research to explore the effect of topology on the brain stability. So we decided to study how the topology affects the balance of resting-state networks. It should be mentioned that our results can be generalized to any signed networks and will be interesting for complex network scientists.

In this study, we hypothesized that the functional signed connections of the brain tend to make hubs mainly by negative links and form a specific non-trivial signed topology that pushes the network toward more stable states with greater numbers of balanced triads. In this regard, we introduced some global measures to clarify the impact of topology on the balance, then investigated them in resting-state signed networks and brought out a mechanism based on the results. Moreover, we studied the emergent behavior of the brain regions in terms of negative degree distributions.

## 2. Results

We summarized the procedure of the study in Fig. 2 We employed the structural balance to assess network stability. The research question was whether the signed network topology (independent variable) can affect the balance (dependent variable) of the resting-state signed network. To answer it, we used a matched-pairs design to compare the balance of actual network (resting-state functional networks) with the balance of null-networks. The null-networks were constructed from shuffled regional activations and had the same node and link size and the same positive to negative link ratio as the actual networks but they had trivial signed link arrangements. Since balance theory works based on signed links and considering the fact that the number of positive and negative links differ in balanced and imbalanced triads (Fig. 1.a), we regarded the equality of positive to negative link ratio to control confounding effects of signed links percentages. Also, we used the matched-pairs design to increase internal validity and decrease the chance of occurring selection bias.

**Figure 2:**
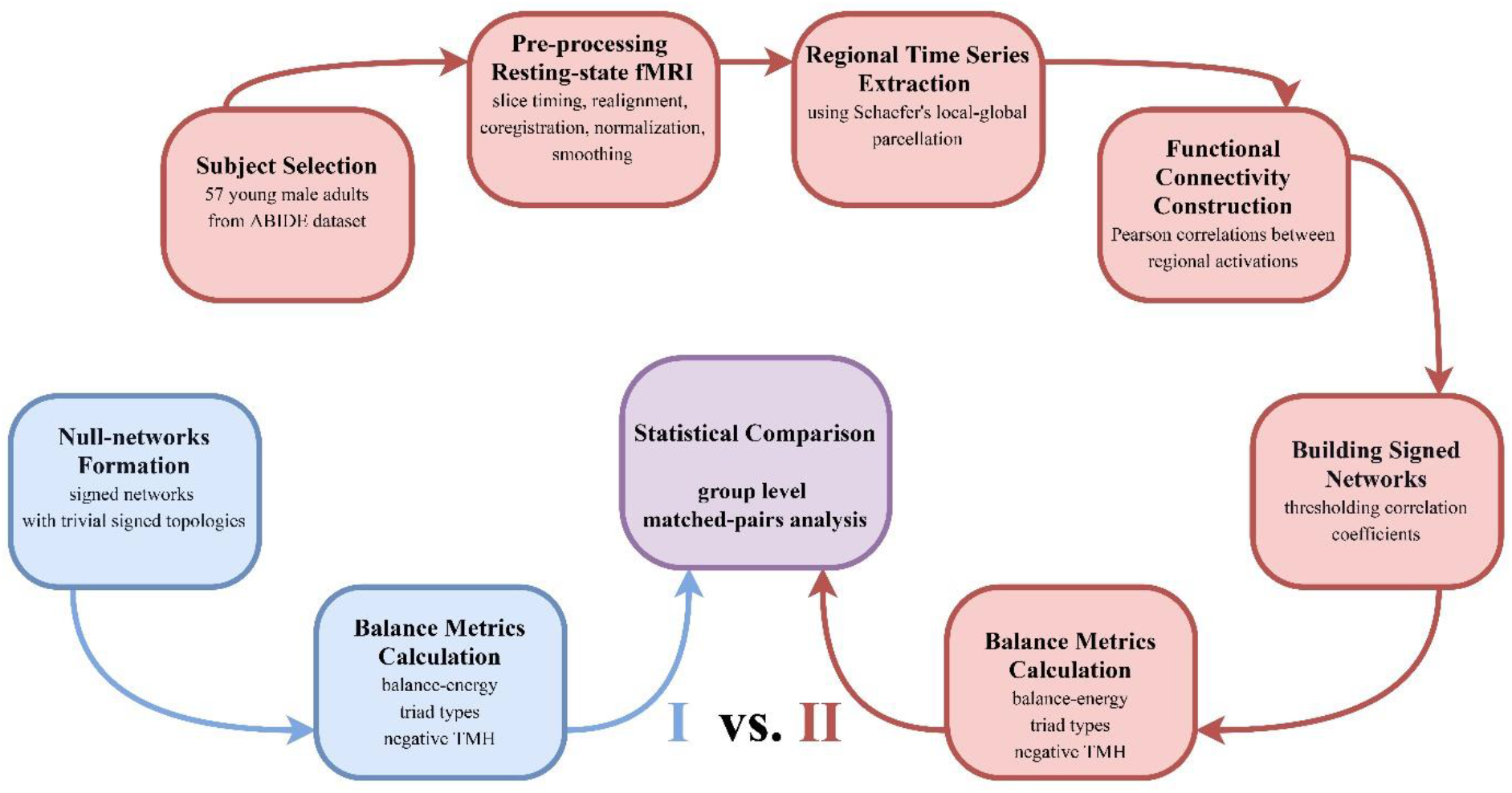
Procedure of the study.

The research procedure is abstractly described as follows. We used publicly shared images of the Autism Brain Imaging Data Exchange (ABIDE) [27] to construct actual networks. Then we only selected 57 right-handed, young male adults to exclude the covariate effect of age, gender, and handedness, and increase the validity of the relationship between independent and dependent variables. After standard preprocessing of functional images, we extracted time-series of regional cerebral cortex activities using Schaefer’s Local-Global parcellation [28]. Then we made functional connectivity for each subject separately and binarized its functional connections to +1 and −1 to formed the resting-states signed network (Fig. 1.b). On the other hand, we created an ensemble of signed networks with trivial topology for each subject whereas their positive to negative link ratio was equivalent to the positive to negative link ratio of the subject’s actual network. Subsequently, we calculated the balance metrics of actual networks and null-networks, compared the balance metrics of the actual network with the ensemble average of balance metrics of the null-networks, then used individual differences for group-level analysis. Since these two types of networks only differ in topology, we imputed the difference between their balances to the topology of the signed links.

### 2.1. Effect of topology on the network balance

To investigate the effect of topology on the network balance, we compared the balance of actual networks with the balance of null-networks. The null-networks were similar to actual networks but have trivial signed link arrangements. As we wanted to perform a paired analysis, we created a set of null-networks matched to each actual network where the matched networks had the same positive to negative link ratios. Afterward, we compared the balance-energy of each actual network and the ensemble average of balance-energies of correspondent null-networks, then we examined the differences for group-level analysis (Fig. 3.a). Since extracted balance metrics of the signed networks did not follow a normal distribution (p-value_Shapiro-Wilkoxon_ < 0.005) and their paired differences around the median were also asymmetric (p-value_Miao-Gel-Gastwirth_ [29] < 0.001) (Supplementary Fig. 1), we decided to use the non-parametric Sign test for the group-level matched-pairs analysis. Fig. 3.a shows that balance-energies of resting-state networks were significantly lower than balance-energies of their paired null-networks (p-value_Sign-test_ < 0.001, effect size = 0.79 (large)). We also compared the percentages of each triad type separately. Fig. 3.b shows that actual networks have significantly a greater number of balanced triads and a little number of imbalanced triads as against null-networks (p-value_Sign-test_ < 0.001, effect size = 0.78 (large)). As actual networks and null-networks differ in signed link arrangement, these results highlight the effect of topology on the balance of the signed networks and indicate that resting-state topology provides balance (stability) for the network.

**Figure 3:**
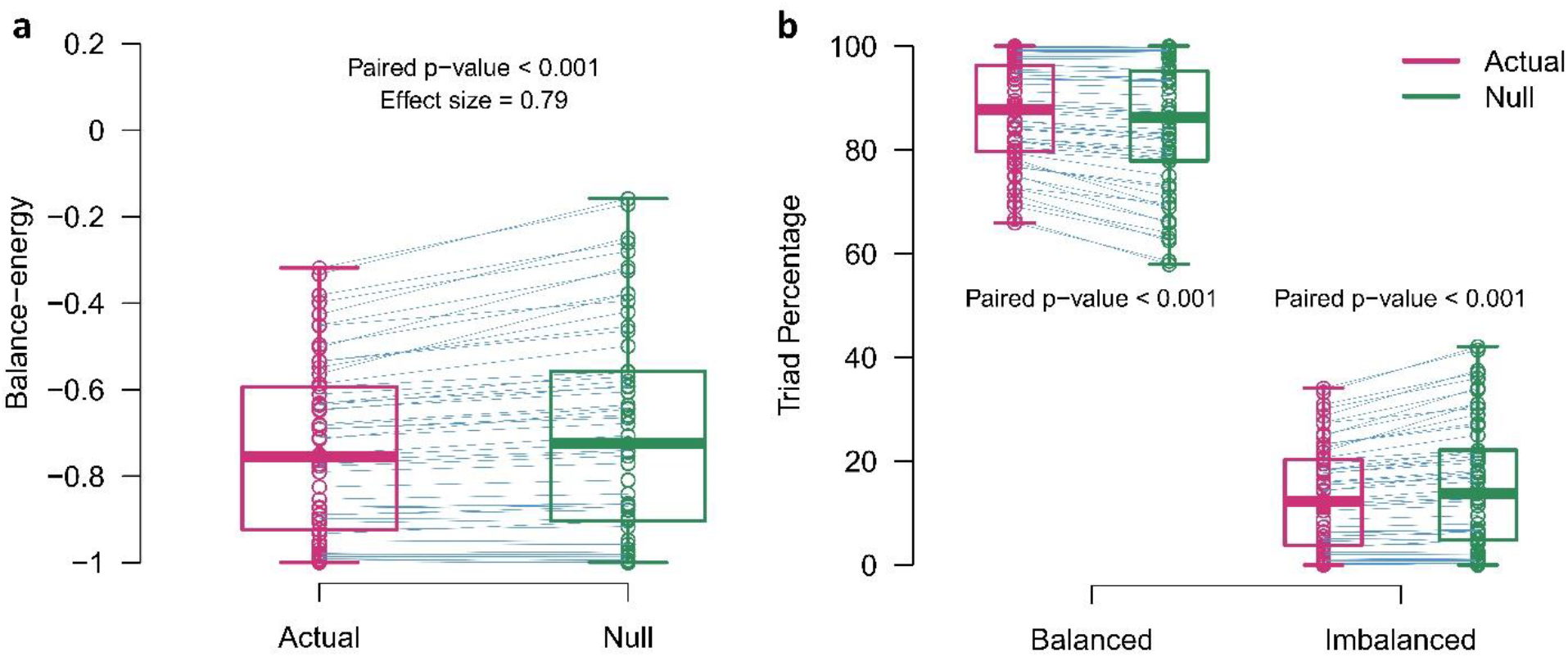
Matched-pairs comparison of balance metrics. **(a)** Balance-energies. **(b)** Percentages of triad types. Violet and green colors denote actual-networks and null-networks, respectively. Circles correspond to the signed networks. The boxes indicate median and interquartile ranges. The blue lines also connect the paired points of the actual and ensemble average of null-networks. P-values and effect sizes are related to the Sign test.

It should be pointed out that we obtained the explained results based on consideration of the fully connected topology. So we applied several thresholds to the functional connections to make partially connected networks then compared the balance-energies to explore the external validity of the results. Supplementary Fig. 2 shows that group-level differences remain significant only for some ranges of the thresholds even after the thresholding process. Nevertheless, the sign of differences switched several times by increasing the threshold. The balance-energy of the resting-state is lower than the balance-energy of the random network from zero to 0.08, higher from 0.14 to 0.22, and again lower from 0.32 to 0.4.

### 2.2. Hubness of functional negative links

In the previous section, we indicated that the actual network and null-networks that were only varied in signed topology had different balance-energies. So we decided to explore the centrality of the network topology. In this regard, we introduced a global hubness measure entitled “Tendency to Make Hub (TMH)” and compared signed TMH of actual networks and null-networks (see method section for further details).

Fig. 4.a shows a matched-pairs comparison between negative TMH scores of actual networks and ensemble average of negative TMH scores of null-networks. Since negative TMHs of networks had non-normal distributions (p-value_Shapiro-Wilkoxon_ < 0.001) and their paired differences distributed symmetry (p-value_Miao-Gel-Gastwirth_ = 0.71) (Supplementary Fig. 3), we used Wilcoxon signed-rank test to explore group-level paired differences. The test revealed that actual networks have larger negative TMH or negative hubness than their correspondent null-networks (p-value_Wilcoxon signed-rank test_ < 0.01, effect size = 0.84 (large)).

**Figure 4:**
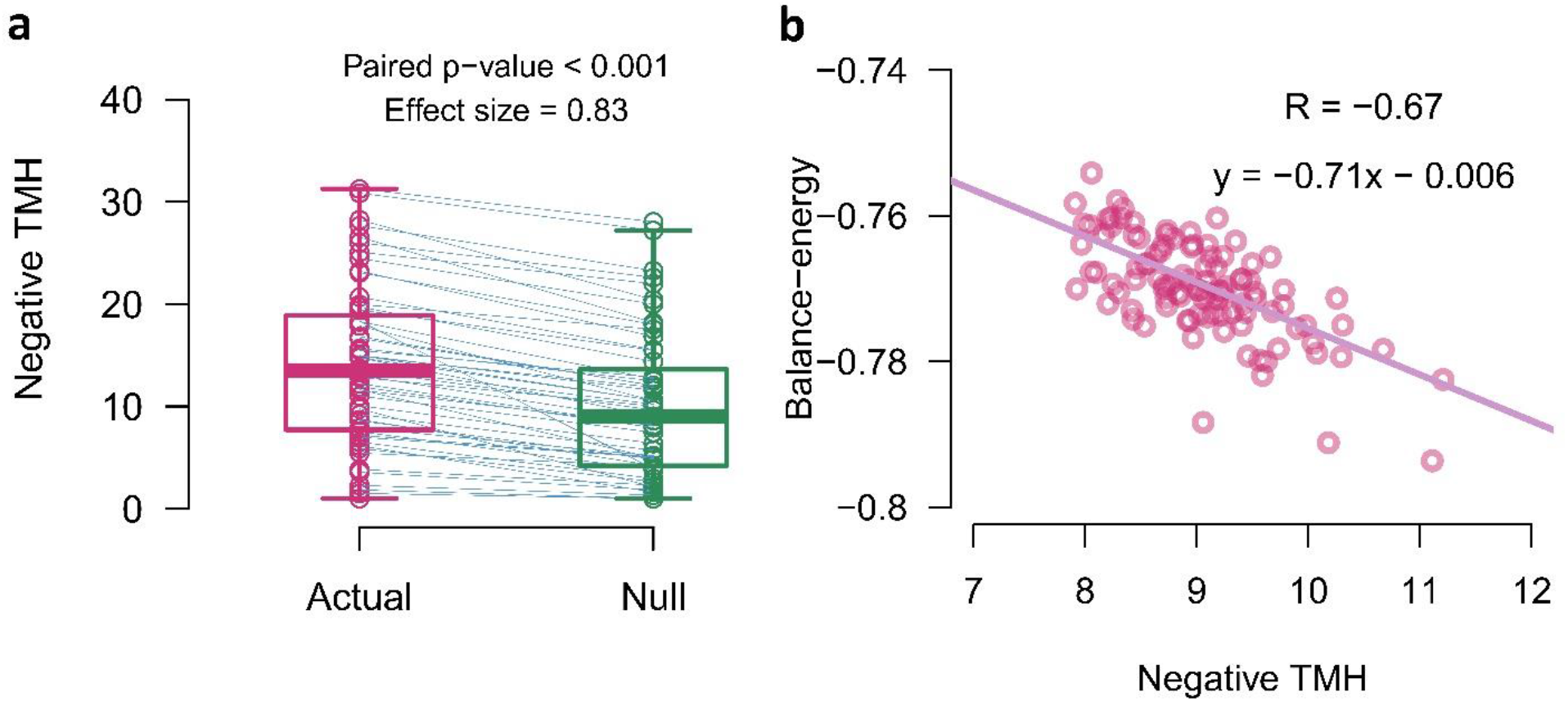
Negative hubness and balance-energy. **(a)** Matched-pairs analysis of negative TMHs between actual networks and their correspondent null-networks. Vertical boxes indicate median and interquartile ranges and blue lines connect paired points. P-value and effect size of Wilcoxon signed-rank test are reported in the figure. **(b)** A negative correlation between the negative TMHs and the balance-energies of simulated networks. Circles denote simulated signed networks and the violet line indicates the linear fitted function to them. Abbreviations: TMH - Tendency to Make Hub; R Pearson correlation coefficient.

### 2.3. Negative hubness and network balance

In the last sections, we indicated that the actual networks have greater negative hubness and lower balance-energy compared to the null-networks. The question arises that is there any relations between signed hubness and network balance. To answer it, we generated some set of fully connected random signed networks (see method section 4.6 for more details) then explored the relation of negative TMH and balance-energy. The generated networks were constrained to have 10 percent negative links but there was no restrictions on signed links arrangements. Fig. 4.b shows a negative correlation between negative TMH and balance-energy (R = −0.67). It means that signed networks with higher signed hubness are more stable.

### 2.4. Description of the effect

In the previous section, we indicated that signed networks with larger negative TMH have lower balance-energies. So, we decided to explore the effect of topology on the network balance in a descriptive manner. Fig. 5.a represents three different topologies of a fully connected signed network with 8 nodes and 4 negative links. The topologies have similar positive to negative link ratios. From left to right, the network topology changes in a way that the negative links tend to gather together and make negative hubs. Mutually, this happens for the positive links moderately, it does not seem significant hence the majority of the links are positive. Increasing negative TMH as a global hubness measure quantifies the hub formation. Also, the balance-energy of the signed network is reduced as a consequence of increasing the number of balanced triads and decreasing the number of imbalanced triads. Additionally, Fig. 5.b schematically describes how the explained mechanism pushes the signed network into a more stable state.

**Figure 5:**
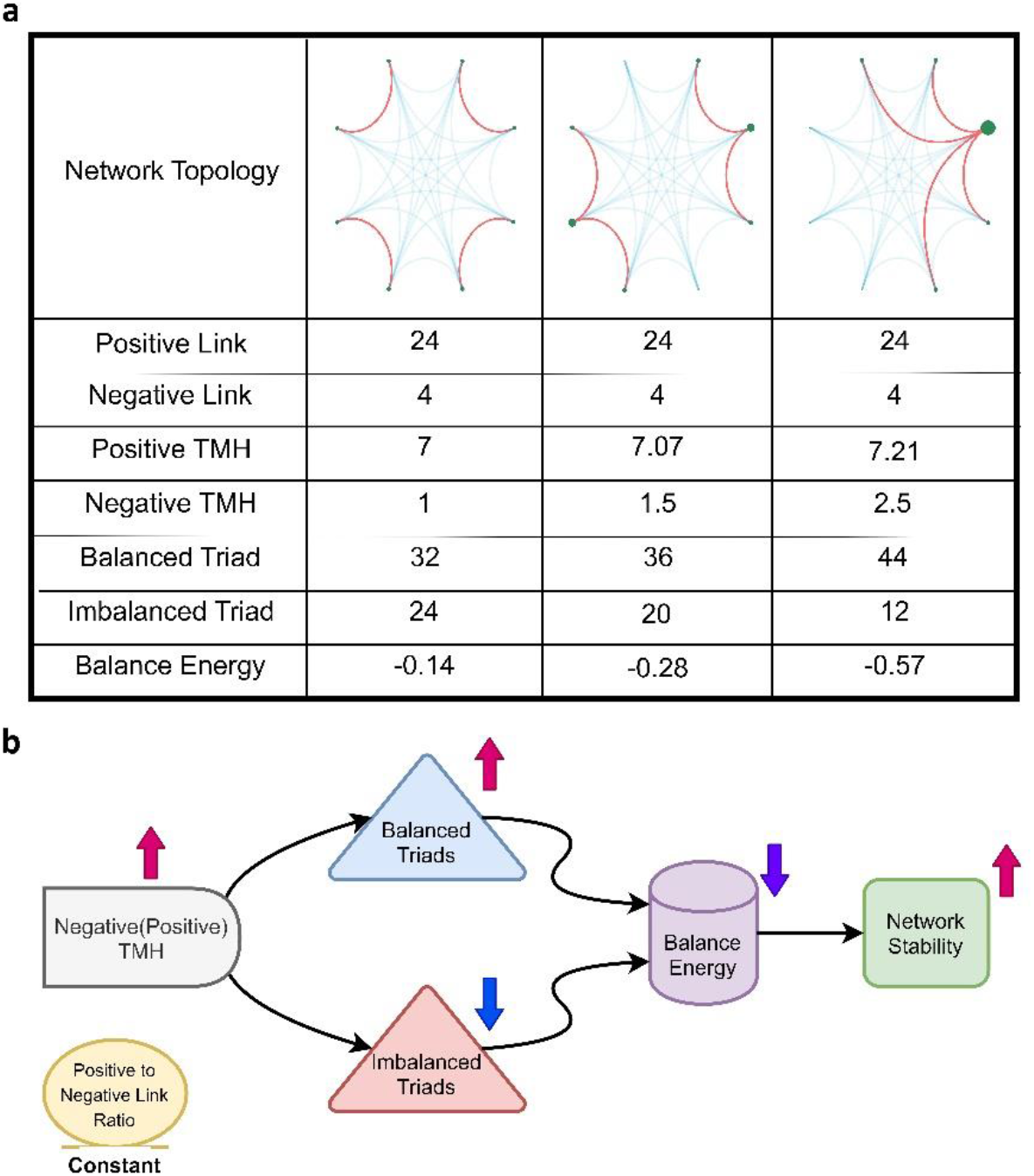
Effect of changing signed topology on the stability of the signed network. **(a)** Descriptive table. Rows demonstrate corresponding metrics of each topology. Positive and negative links of networks are indicated with blue and red curved lines, respectively. The size of green circles displays negative degrees of nodes. **(b)** Schematic diagram. Abbreviation: TMH – Tendency to Make Hub.

### 2.5. Collective properties of negative links

Although minor of resting-state functional connections are anti-synchronous (negatively correlated) (Supplementary Fig. 4), their collective behavior has the most magnificent effect on the network balance. Fig. 5 also confirms this fact. Therefore we decided to explore the collective behavior of resting-state negative links.

So we decided to explore the distribution of nodal negative degrees. Fig. 6.a shows the logarithm of the negative degree distribution of all the subjects. Linear functionality of the logarithmic distribution denotes that negative degrees distribute exponentially. We used the Maximum Likelihood Estimation to assess the rate parameter of the exponential distribution (λ_resting-state network_ = 0.14). The figure also represents the negative degree distribution of the null-networks. The difference between the right tail of distributions highlights the presence of negative hubs in actual networks despite null-networks.

**Figure 6:**
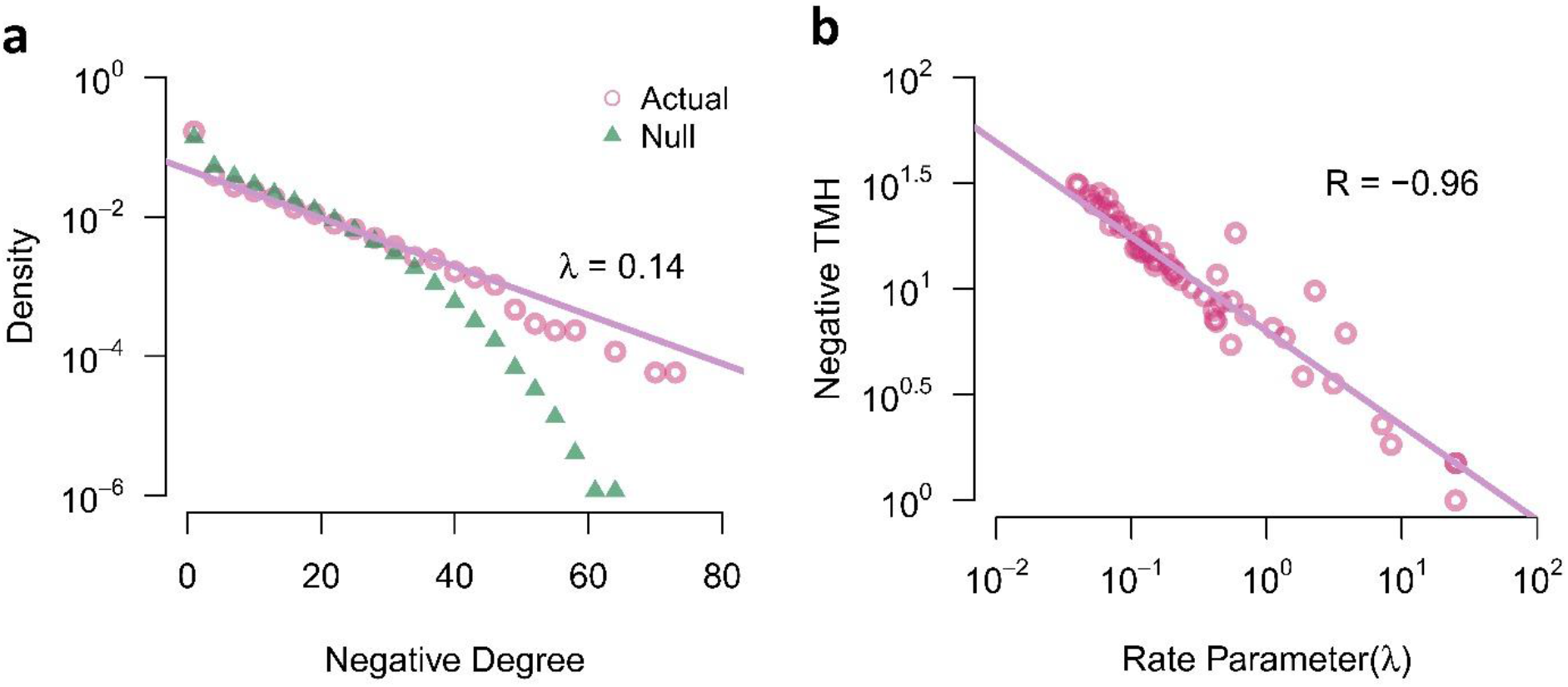
Collective behavior of resting-state negative functions. **(a)** Semi-logarithmic negative degree distribution of actual networks and null-networks. **(b)** A negative correlation between the logarithm of negative TMH and the logarithm of the rate parameter of negative degree distribution. Each circle corresponds to the signed network of a subject. Abbreviations: TMH – Tendency to Make Hub; R Pearson correlation coefficient; λ – rate parameter of the exponential degree distribution.

In fact, when we study exponential distributions in logarithmic form, the sharpness of degree distribution is associated with the hubness of the network. Therefore, we explored the relation between the negative TMH of the actual networks and the rate parameters of the negative degree distribution of the actual networks. We found a negative correlation between the logarithm of rate parameters and the logarithm of negative TMHs (R = −0.96) (Fig. 6.b). This result indicated that actual networks with lower rate parameters and lower sharpness of distributions have higher negative hubness.

### 2.6. Emergence of brain regions

Since investigating the collective behavior of functional negative links brought out interesting results, we decided to explore the existence of emergent behavior between brain regions in terms of negative degree distribution. Emergence is a property of complex systems that occurs when individuals behave in a different manner as compared to the whole. Each brain region has a distribution that contains negative degrees of subjects. We compared the negative degree distribution of each region with the negative degree distribution of the whole brain using the Kolmogorov-Smirnov test. Statistical tests indicated that negative degrees of some cortical regions differently distribute than the whole. Fig. 7 represents significantly different ROIs after a multiple comparison correction using the False Discovery Rate (FDR) method (corrected p-value < 0.05). The significant ROIs locate at right and left precuneus/posterior cingulate of default mode network, left frontal operculum of the insula, left median of ventral attention network, and right frontoparietal control network. Please see Supplementary Table 1 for details. Supplementary Fig. 5 also displays the results without multiple comparison correction.

**Figure 7:**
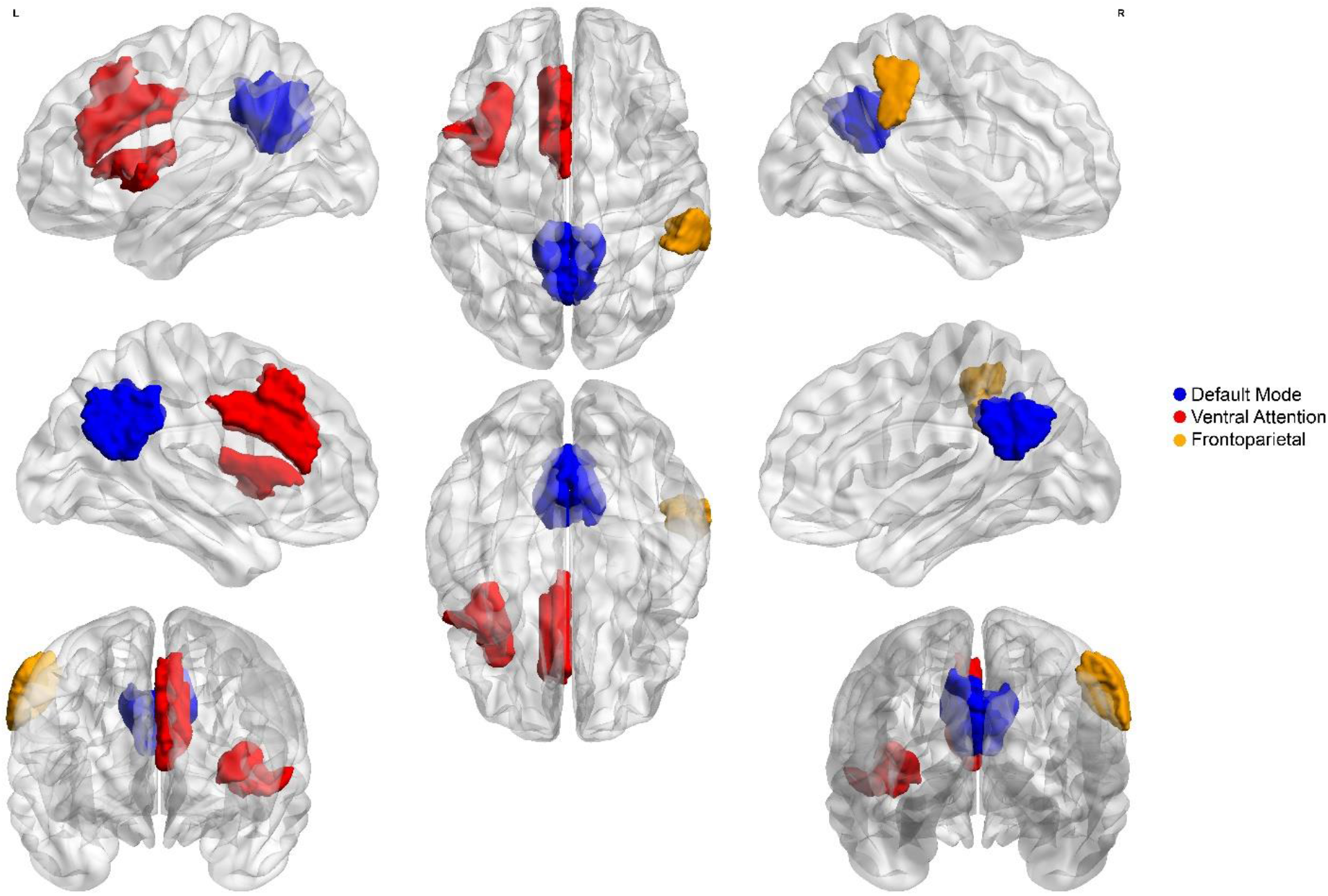
Emergence property of regions of interest. Colored portions of the brain images show regions that their negative degree distributions significantly differ from the whole-brain negative degree distribution (multiple comparisons corrected). Various colors denote that each region belongs to which large-scale cortical networks. The brain maps were created using BrianNet Viewer toolbox [30] of the Matlab (http://www.mathworks.com/products/matlab/).

## 3. Discussion

In brief, we indicated that functional negative links of the resting-state network formed an especial topology to push the network toward more balance (stable) states. In this regard, we compared actual networks and null-networks with trivial topologies and the same positive to negative link ratio. We showed that the actual networks have lower balance-energy as compared to the null-networks. We also introduced a global measure of hubness entitled Tendency to Make Hub (TMH) and showed that the actual networks have higher negative TMH against the null-networks. According to the results, we brought out a mechanism to describe how signed hubness causes network stability. Since we found that negative links of the brain have an influential role in network stability, we decided to explore the collective behavior of them. We discovered functional negative links of resting-state networks distribute exponentially and the rate parameter of the distribution negatively correlates with the negative TMH. Also, we affirmed the emergence of brain regions in terms of negative degree distribution where some brain regions do not follow the behavior of the whole-brain.

### 3.1. Definition of brain energy

In recent years, some studies have defined the energy of neuroimaging data at a particular moment using brain regional activations at that moment multiplied by long-term coactivations between regions [31–35]. Since they used momentary activations and considering the fact that fMRI signals have low signal-to-noise-ratio, the validity of calculated energies may be questionable. Since we used long-term coactivation in calculating energy we kept away from this issue.

### 3.2. Low energy resting-state signed network

Fig. 3.a showed that the balance-energy of the resting-state network is negative and it posits near the absolute stable state. It not only depends on the topology but also is a consequence of positive and negative link appearance. Exploring this issue may disclose why most brain regional interactions are synchronous, not anti-synchronous.

### 3.3. Null-network formation

As we explained, we hypothesized that the special topology of signed links affects the stability of the resting-state network. To test the hypothesis, we compared the balance of actual networks with the balance of null-networks. Considering the validity of statistical inference highly depends on null-network selection, we innovated a null-network formation procedure to control confounding variables. In addition to equality of node size and link size, null-networks had positive to negative link ratios similar to actual networks and they were extracted from shuffled regional brain activations to remain signal distributions consistent. As the null-networks only varied in signed link arrangement, we attributed the difference between the balance of actual networks and the balance of null-networks to the topology.

Since two balanced triads totally have greater numbers of positive links and two imbalanced triads totally have greater numbers of negative links (Fig. 1.a), we had to regard the condition of equality of positive to negative link ratio to control confounding factor of sign link presence. Although there are various approaches to construct null-networks [36], since none of them focused on the signed networks and balance theory works based on signed links interactions, we decided to create our procedure to form proper null-networks.

In our null-network creation procedure, we used signals to construct null-networks, not signed links. It enabled us to apply adjusting signals and form null-networks with desired positive to negative link ratios.

It is also necessary to point out another benefit of using signals. As we investigated triadic interactions of brain regions and considering the fact that dual relations of a triadic relation in the brain are not independent, generation of null-networks from signed links disregarding constraints on triadic interrelations could be problematic. In other words, a correlation between region A and region B and a correlation between region A and region C restrict correlation between B and C which was not considered when we only assigned signed links to a network.

### 3.4. Thresholding functional connections

Network neuroscientists usually apply thresholds on the correlation coefficients to remove specious connections and magnify key topological features of the networks [37]. Since the impact of positive links and negative links on the balance of the resting-state network were not equal, we claimed that the negative weak correlations should not be ignored when we study network balance, so we decided to consider signed networks fully-connected. Nevertheless, we employed a thresholding process on the connections and obtained significant differences between the balance-energies of actual networks and balance-energies of null-networks in some range of thresholds (Supplementary Fig. 2). We also observed that sign of group-level differences switches by increasing the threshold. It reveals the importance of further investigation in the effect of the thresholding process on the signed network balance.

### 3.5. Effect of topology on the balance

After more than half a century since the appearance of Heider’s balance theory, there is no research work to clearly describe the origin of the balance in the signed networks. Typically, network balance is determined by counting the triad types. It seems more an observative fact than a causal effect. So, it is not still clear what emerges the triad types? In this study, we concluded that the network topology plays an influential role in triad type appearance. We described that gathering sign links around the nodes and making signed hubs increases the number of balanced triads and decreases the number of imbalanced triads (Fig. 4.b and Fig. 5).

### 3.6. Exponential negative degree distribution

Network neuroscientists usually regarded both positive functional connections and negative functional connections as links and construct brain network without considering signed of functional connections [37]. In addition, sometimes they ignored negative connections due to the difficulties in the interpretation and justification of anti-correlated activations [38]. But in this study, we highlighted the role of functional negative links. In accordance with the work of Valerio Ciotti et al. [39], we extracted positive and negative subnetworks and explored them separately. We studied the collective behavior of resting-state functional negative links and found out that they distribute exponentially (Fig. 6.a). Whereas previous works indicated a power-law distribution without any consideration on the sign of the links [40–43], we assigned a new attribute to the collective behavior of functional negative links and emphasized the study on the anti-synchrony in the brain functional network.

### 3.7. Emergence of the resting-state “signed” network

Ordinarily, brain networks have been considered unsigned. They emerge complex properties such as small-worldness that facilitate the integration of information and segregate brain functions [24]. Based on our knowledge, there is no outstanding research that regards the brain as a signed network. Since we considered brain network signed to determine network balance, we think it is necessary to explore the complexity of resting-state signed network regarding signed links. So we explored the emergence of brain regions in terms of negative degree distribution. Emergent property explained that individuals may work differently from the whole in a complex system [44]. We found that the negative degree distribution of some brain regions significantly differed from the degree distribution of the whole brain. It is in agreement with the emergent behavior of a complex system.

### 3.8. Metastability of resting-state signed networks

Metastability refers to a stable state other than the state of least energy (ground state). A metastable state has a shorter lifetime than the ground state and is the place that transition to other states is likely to occur [45]. Some studies indicated metastability of the brain regarding brain dynamics [46–47], but we observed it in a static manner. Fig. 3.a indicates that resting-state functional networks inhabit at a low energy level near the absolutely stable state, we claimed that is a metastable state.

### 3.9. Limitations and considerations

We had to select our imaging data from various studies. Although most of the scan parameters including repetition time (TR) and the length of scanning were similar, other scanning parameters such as echo time (TE) might vary.

It has been shown that applying global signal regression may induce anti-synchronous activities and increase the number of negative links [38, 48–49]. Considering the recent neuroimaging studies that suggest not remove the global signal [50–53], we did not perform global signal regression in the preprocessing steps. Nevertheless, we performed our analysis with global signal regression as well. The results were still valid and are presented in Supplementary Fig. 6.

The main consideration of the current study is about the semantic of balance-energy. In this paper, the energy term does not refer to physical energy which roots in biological metabolic actions; but, it is a metaphor to explain the global behavior of a signed network. It should be considered that, to the best of our knowledge, there was no application of balance theory in neuroscience so far.

### 3.10. Conclusion and future directions

In summary, we concluded that negative functional connections of the resting-state brain network make hubs to form an especially signed network topology and push the network toward more balance states with lower balance-energies. Negative links play an important role in the stability of resting-state networks, their collective behavior is not trivial, can expose the complexity of brain regions. So further researches are needed to investigate them.

Nevertheless, we only investigated the balance of resting-state networks in young adults. However, more investigation during the development and degeneration of the mature brain would be required in future works. Also, stability and phase transition of task-dependent and dynamic functional networks could be traced by the balance theory. We also expect that the balance-energy of functional brain networks in neurodevelopmental disorders such as autism falls into a jammed-state (metastable states) that restricts their behavioral performance. Lastly, we hope the introduced topological basis of balance could bring new solutions for escaping from that jammed-states.

## 4. Method

### 4.1. Neuroimaging data

We selected 70 healthy male adults from 2226 available subjects of the ABIDE repository [27]. Fig. 8 shows the subject selection process. All of the subjects were right-handed and aged from 18 to 31. Each participant underwent a T1-weighted structural MRI and a resting-state fMRI scan in the same session. The high-resolution T1-weighted structural MRI was acquired using magnetization-prepared rapid. The resting-state fMRI was also acquired using a single-shot EPI sequence during eyes-open task-free condition with a repetition time (TR) of 2 seconds. We reported the demographics of the subjects and scanning parameters of the selected dataset in Supplementary Table 2. Finally, we eliminated some cases with destructive artifacts (movement parameters more than one voxel size) to derive 57 subjects (age = 24+-4).

**Figure 8:**
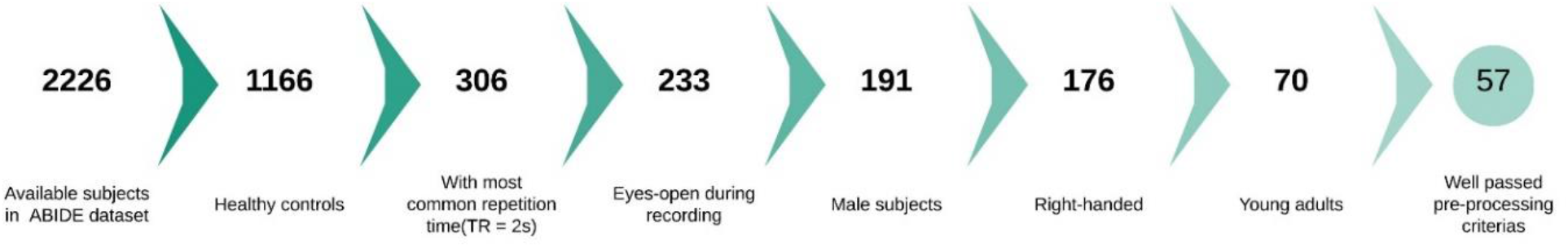
Subject selection procedure.

### 4.2. Pre-processing of resting-state functional images

We used a standard preprocessing pipeline that utilizes the FMRIB software library v5.0 (FSL: http://www.fmrib.ox.ac.uk/fsl) [54] and Analysis of Functional NeuroImages environment (AFNI: http://afni.nimh.nih.gov/afni) [55]. We deobliqued all of the structural images to FSL friendly spaces and extracted the brain. Then, we discarded the first five volumes of resting-state fMRI images to ensure the magnetization stability. Subsequently, we carried out slice time correction for interleaved acquisitions using Fourier interpolation. After that, we registered 3D volumes of functional images to their corresponding high-resolution structural images of native space using the least square algorithm (3 translational and 3 rotational variables were optimized). Then, we interpolated outlier time points using a continuous transformation function and normalized each voxel to the average of its activities. We then applied spatial smoothing using a Gaussian kernel function with Full width at Half Maximum (FWHM) equal to 5mm, and we performed temporal bandpass filtering (0.009-0.01 Hz) on the functional images. To prepare for group-level analysis, we nonlinearly registered 3D images to MNI152 standard space using optimizing 12 variables which were related to translation, rotation, scaling, and shearing. Then, we regressed out confounds of motion (3 translational and 3 rotational parameters), white matter (WM) and cerebral spinal fluid (CSF). Finally, we checked whether movement parameters of the functional images to be less than one voxel size and visually inspected the quality of brain extraction and segmentation in structural images. In this regard, 13 out of 70 subjects could not pass the imaging criteria, so we removed them from further analysis. We have performed and checked the above-mentioned procedure in our previous works as well [26, 56, 57].

### 4.3. Activity pattern of the brain regions

We used MATLAB software to extract time courses of the brain regional activities. We chose Schaefer’s Local-Global atlas [28] to parcellate the cerebral cortex into 100 homogenous regions of interest (ROIs). In this atlas, each parcel is located in one of Thomas Yeo’s canonical networks [58]. To extract the activity pattern of a region, we multiplied the binary mask of that ROI to 3D images and considered the average of BOLD signals in each 3D image as activity of ROI at that time point. In this way, we drew out 100 time-series from the fMRI image of each subject. The number of recorded 3D images (volumes) in the functional imaging determined the length of the time series. Considering that we used functional images from various imaging-sites with different numbers of recorded volumes, we peaked up the extracted times series equaled to the shortest ones. We carried out the equalization process because the length of time courses might affect the temporal correlations between regions and connectivity matrices.

### 4.4. Resting-state signed networks

Functional connectivity represents temporal synchrony between brain regional activations. The synchrony between two regions also is defined as the Pearson’s correlation coefficients of their temporal activations. In this way, activity patterns of the brain regions may be positively correlated (synchronous) or negatively correlated (anti-synchronous). The correlations are considered as elements of the functional connectivity matrix. Finally, we binarized the elements of connectivity matrices to +1 and −1 by considering the sign of correlation coefficients and built a resting-state signed network for each subject (Fig. 1.b).

### 4.5. Structural balance

Structural balance theory studies collective behavior and stability of signed networks based on triadic associations between entities. The theory is rooted in Fritz Heider’s researches on attitude change then applied to interpersonal relations [59, 18]. According to the theory, if you are a friend of your friend’s friend or an enemy of your friend’s enemy, they are trivial and your triadic relation is balanced (stable); otherwise, if you are an enemy of your friend’s friend or an enemy of your enemy’s enemy, your triadic relation is imbalanced (unstable) [19]. These 4 triadic relations can be modeled using signed links (Fig. 1a). It should be noted that an imbalanced triad is considered a frustrating condition and endures tension to change their links to become balanced.

We can explore the balance of a signed network where there are lots of triad types [21]. More balance networks have a larger number of balanced triads and less number of imbalanced triads. So, the balance-energy of a signed network defines as the difference between the number of balanced triads and the number of imbalanced triads [22] as follows:

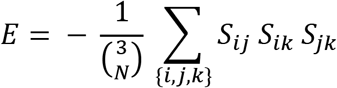

The summation is performed on the possible triads of the network. *S_ij_* denotes a signed connection between node i and node j. *S_ij_ S_ik_ S_jk_* also represents the multiplication of edge values in a triad where — *S_ij_ S_ik_ S_jk_* is balance-energy of the triad and equal to either −1 or +1 for balanced or imbalanced triads, respectively. Minus sign behind the multiplication helps to better understand the equation from the physical energy perspective. 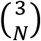 also denotes the number of 3-combination from N elements which is equal to the number of possible triads in an N-node network. This term plays the role of normalization and confines balance-energy between −1 and +1.

The stability of a signed network is associated with the minimization of its balance-energy. Actually, the most stable state when all triadic relations are balanced has the lowest balance-energy equal to −1, and the most unstable state when all triadic relations are imbalanced has the highest balance-energy equal to +1.

### 4.6. Creating null-networks

We innovated a procedure to form null-networks to compare their balance metrics with the balance metrics of signed resting-state networks. We provided a set of null-networks matched to each actual network. The null-networks had the same number of nodes and links and positive to negative link ratio as the actual network but differed in signed link topology. We created a null-network corresponded to an actual network as follows:

At first, we shuffled time points of regional brain activations (which make the actual network) and built new signals. These shuffled signals had similar lengths and distributions to the actual regional activations. It is clear that if we wanted to construct connectivity from these shuffled signals, the connectivity matrix had an equal number of positive and negative connections. It may conflict with the nature of brain functional connectivity where most of the connections are positive and is not appropriate for testing our hypothesis. Therefore, we decided to add a random signal to all of the shuffled signals before calculating the connectivity. The random signal had a normal distribution with “a mean and a variance” equal to “the mean and the variance of regional activations”. In this way, we could increase shared information between the shuffled signals and increase the number of positive connections (positive links). Actually, we multiplied the random signal by an adjusting coefficient then added it to the shuffled signals. So, by changing the coefficient of summation we could adjust the number of positive and negative connections and make it similar to the actual brain network. In this way, we equalized the positive to negative link ratio of the null-network and the actual network. Fig. 9 schematically describes the procedure.

**Figure 9:**
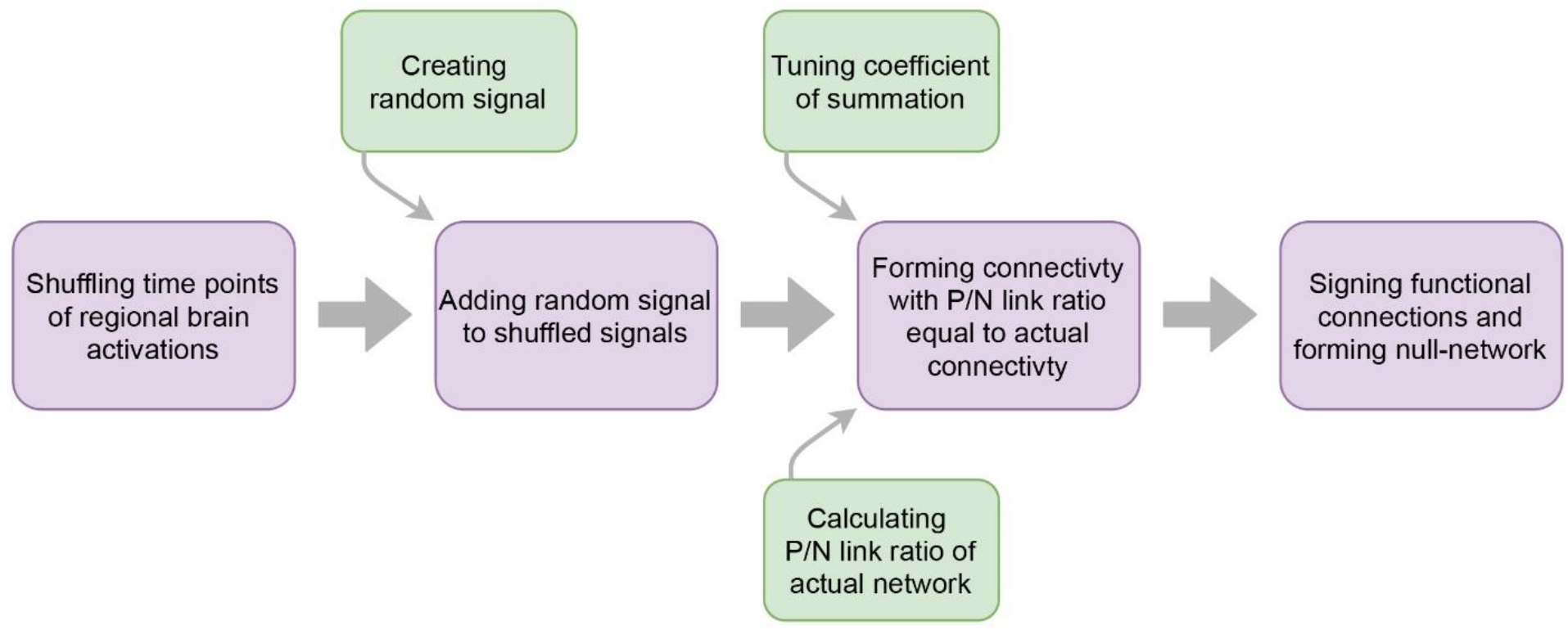
Procedure of null-network formation.

In general, we produced 1000 null-networks corresponded to each actual network, then compared the balance metrics of each actual network with the average ensemble of the balance metrics of corresponded null-networks, and finally used the calculated differences for group-level analysis (Fig. 3 and Fig 4.a). We also explained why we chose this procedure to construct null-networks in the discussion, at “null-network formation” section.

### 4.7. Tendency to Make Hub

Measures of the complex networks are either local or global. The global measures assess the collective properties of the network and the local measures explain features of the network elements [60]. A hub is a local feature of a network that is assigned to the nodes with extremely connected links [61]. In this study, we introduced a novel global hubness measure and named it “Tendency to Make Hub (TMH)” to quantify the strength of link gathering around the nodes in the network from a global perspective. The TMH is defined for a networks as follows:

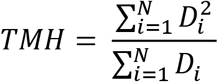

where *D_i_* denotes the degree of an individual node, and N represents the total number of nodes in the network. The TMH takes the value of 1 or above since the degrees of the nodes have values of 1 or above. In fact, the TMH has a higher value when the links have more tendency to assemble around the nodes and make more hubs. In this study, we were interested in the assemblage of the negative links; so, we defined sign-dependent TMH as follows:

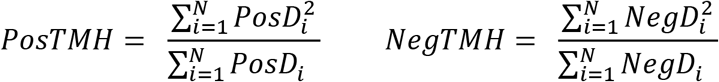

where “positive TMH (*PosTMH*)” demonstrates the tendency to make hub with positive links and “negative TMH (*NegTMH*)” indicates the tendency to make hub with negative links. In this regard, we defined the number of connected positive links to a node as “positive degree (*PosD*)” and the number of connected negative links to a node as “negative degree (*NegD*)” of that node. “Signed degree distribution” also refers to the distribution of positive degrees or negative degrees.

### 4.8. Statistical analysis

To perform a matched-pairs group analysis between balance-energies of resting-state networks and ensemble averages of balance-energies of random topology networks (Fig. 3.a), initially, we tested group-level normality of the balance-energies using the Shapiro-Wilcoxon test. In this way, we found that the distribution of the balance-energies was far from normal and this finding led us to use non-parametric statistics. Then, we employed the Miao-Gel-Gastwirth test [29] to check the symmetry of paired differences distribution. The results showed that the differences were asymmetric, therefore, we utilized the Sign test which is a non-parametric statistical method for matched group-level analysis when the paired differences distribute asymmetrically. We also carried out this procedure for the percentages of the triad types (Fig. 3.b).

Also, to compare the negative TMHs of actual networks and negative TMHs of null-networks (Fig. 4.a), at first we applied Shapiro-Wilcoxon and Miao-Gel-Gastwirth tests. The Shapiro-Wilcoxon test showed the non-normality of negative TMHs as similar to the balance-energies, but Miao-Gel-Gastwirth could not reject the null hypothesis of symmetry of the paired-differences distribution. In this situation, we had two choices for the non-parametric matched-pairs analysis, the Sign test and the Wilcoxon signed-rank test. So we chose the Wilcoxon signed-rank that provide greater statistical power.

We also applied Maximum Likelihood Estimation using Nelder-Mead algorithm [62] to fit an exponential model to negative degree distribution of actual networks and extract correspondent rate parameter (Fig. 6).

Moreover, to explore the emergent behavior of brain regions from the aspect of negative degree distributions (Fig. 7), we compared the negative degree distribution of each ROIs to the negative degree distribution of the whole brain using Kolmogorov-Smirnov test. Subsequently, we corrected p-values from false positives caused by multiple comparisons using the False Discovery Rate (FDR) method developed by Benjamini & Hochberg algorithm [63].

We used the R software [64] and some of their packages [65–69] for statistical analysis and creating graphical figures. We also created brain images (Fig. 7 and Supplementary Fig. 5) using BrianNet Viewer toolbox [30] and drew diagrams (Fig. 2, Fig. 9, and Supplementary Fig. 7) by Draw.io free online diagram editor [70].

In addition, we shared our codes on “https://github.com/majidsaberi/NegLinkTopoBalance“. So everyone can replicate and develop our work.

## 6. Acknowledgments

We would like to thank autism brain imaging data exchange (ABIDE) for generously sharing the data.

## Contributions

M.S, R.K, G.J and A.K designed the study. R.K, G.J, A.K, B.M supervised the research. M.S and R.K analyzed the data and wrote the manuscript. M.S drew the figures. G.J, A.K and B.M revised the manuscript.

## Corresponding author

Reza Khosrowabadi

