## Supplementary material for "Topological impact of negative links on the stability of resting-state brain network"

***Corresponding author:** Reza Khosrowabadi

**Supplementary figures:**

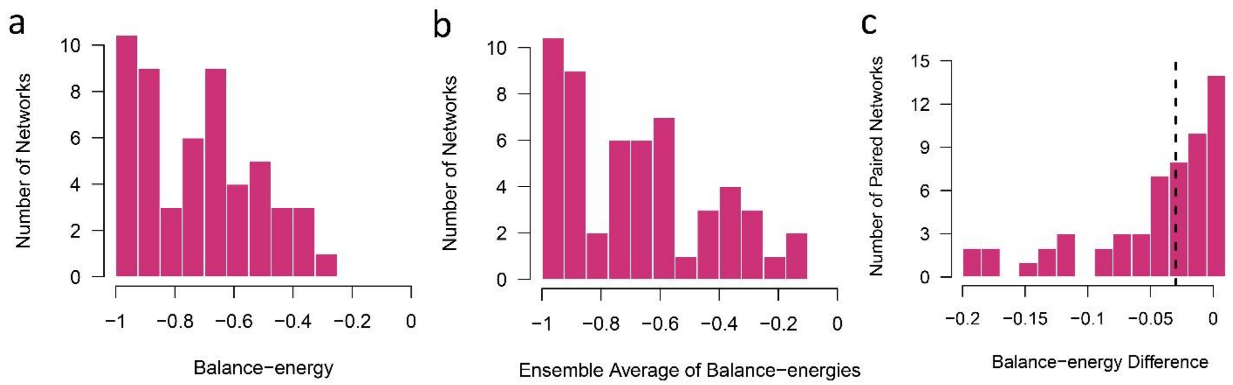


**Figure S1: Histogram of balance-energies (a)** Balance-energies of actual networks. **(b)** The ensemble averages of balance-energies of null-networks. **(c)** Differentiations of balance-energy between actual networks and correspondent ensemble average of null-networks.

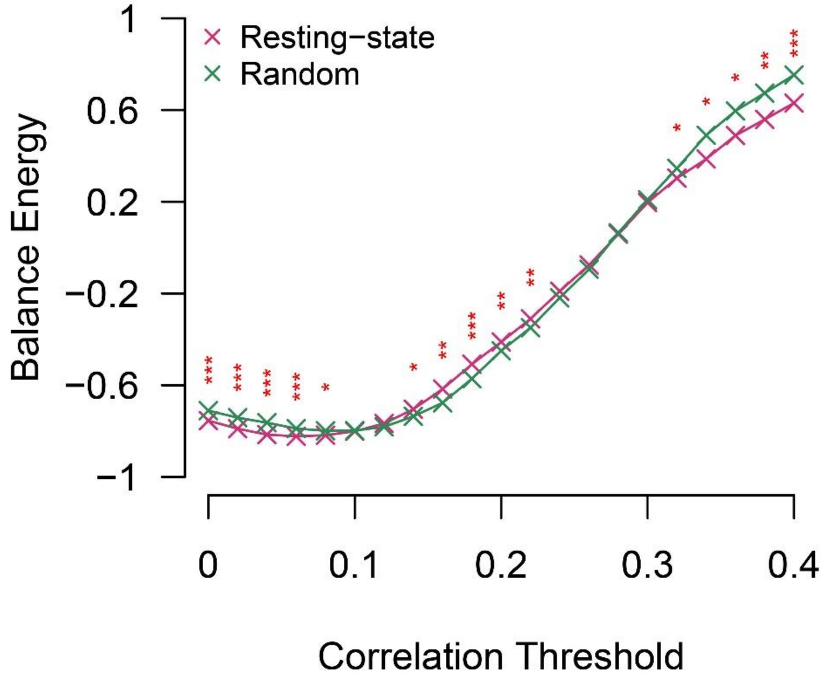


**Figure S2: Considering the threshold on the links before matched-pairs comparison of balance-energies between the actual networks and correspondent null-networks.**

*: Significance of p-value <0.05

**: Significance of p-value <0.01

***: Significance of p-value <0.001

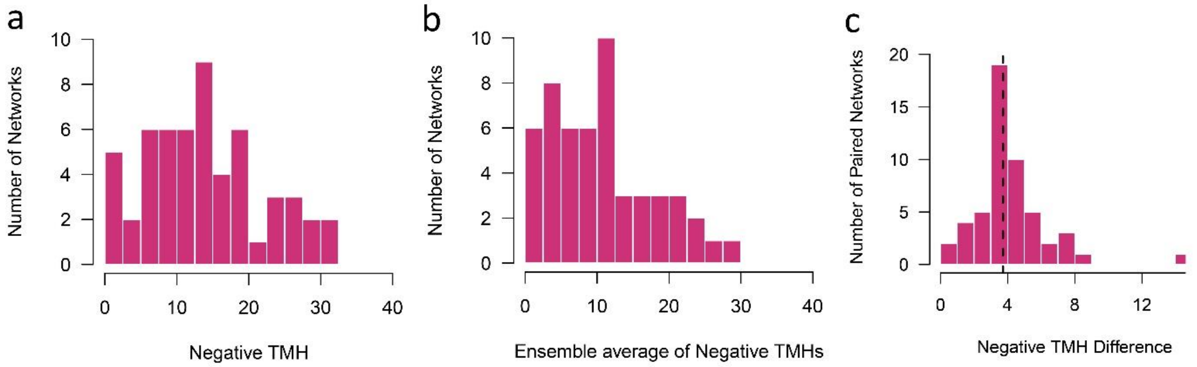


**Figure S3: Histogram of Negative TMHs (a)** Negative TMHs of actual networks. **(b)** The ensemble averages of negative TMHs of null-networks. **(c)** Differentiations of negative TMHs between actual networks and correspondent ensemble average of null-networks.

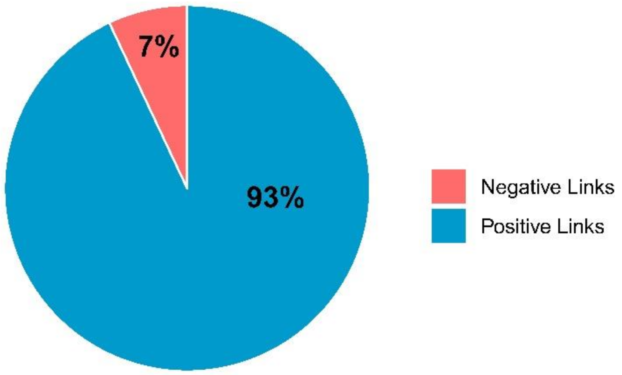


**Figure S4: Signed links Percentages of resting-state networks.**

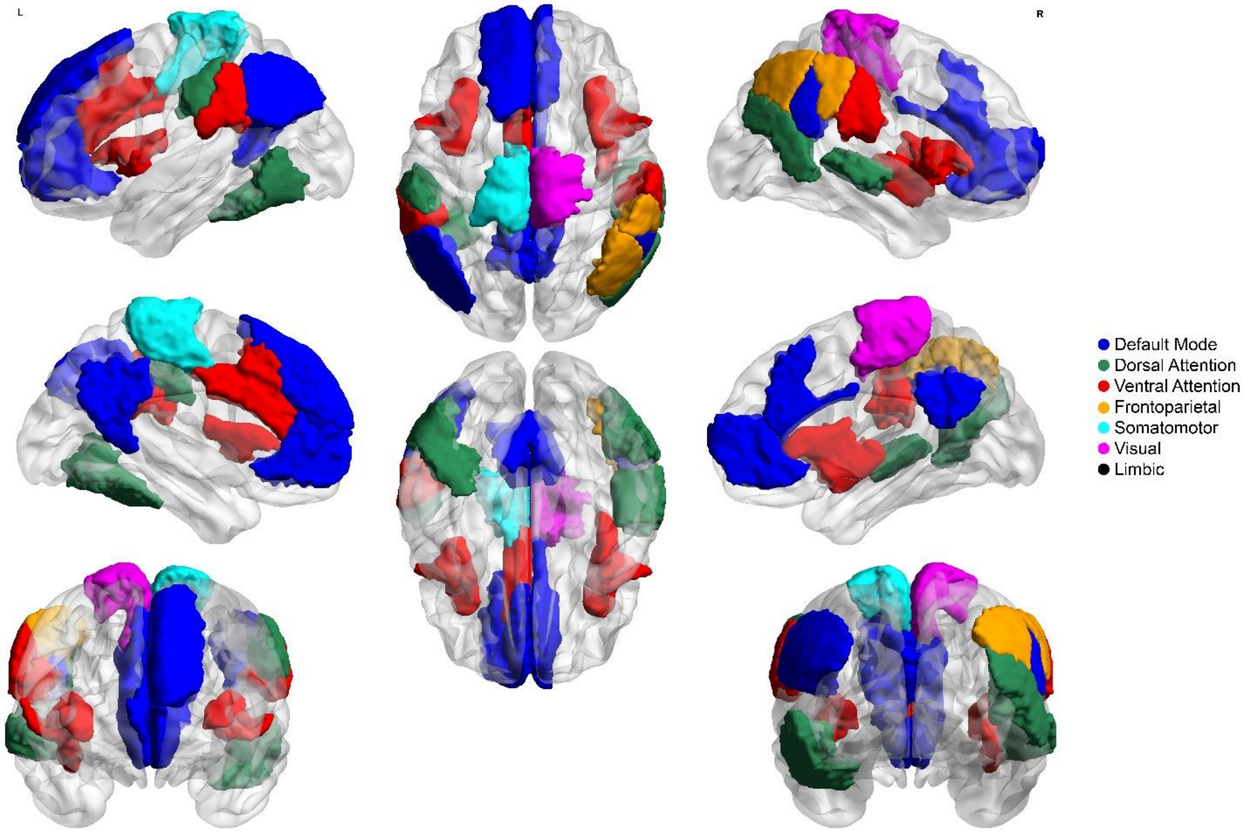


**Figure S5: Emergent property of regions of interests.** Colored parts show regions that their negative degree distributions significantly differ from the whole-brain negative degree distribution (without multiple comparison correction). Various colors indicate significant regions belong to which large-scale cortical networks. The brain maps were created using BrianNet Viewer toolbox of the Matlab (http://www.mathworks.com/products/matlab/)

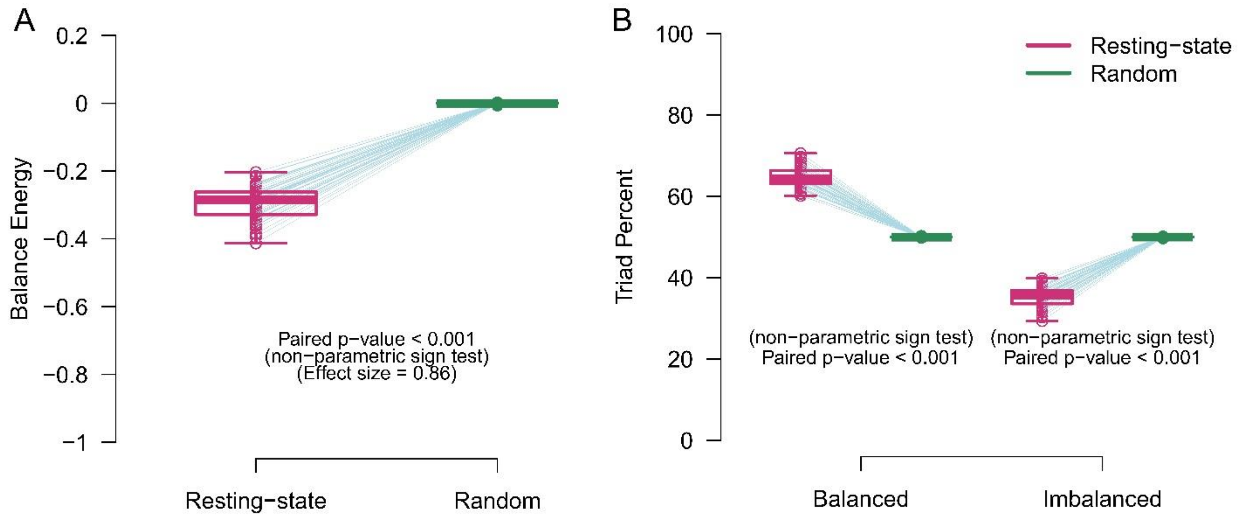


**Figure S6: Matched-pairs comparison of balance metrics when we applied Global Signal Regression. (a)** Balance-energies. **(b)** Triad types. Circles correspond to the signed networks. The boxes indicate median and interquartile ranges. The blue lines also connect the paired points of the actual networks and ensemble averages of null-networks.

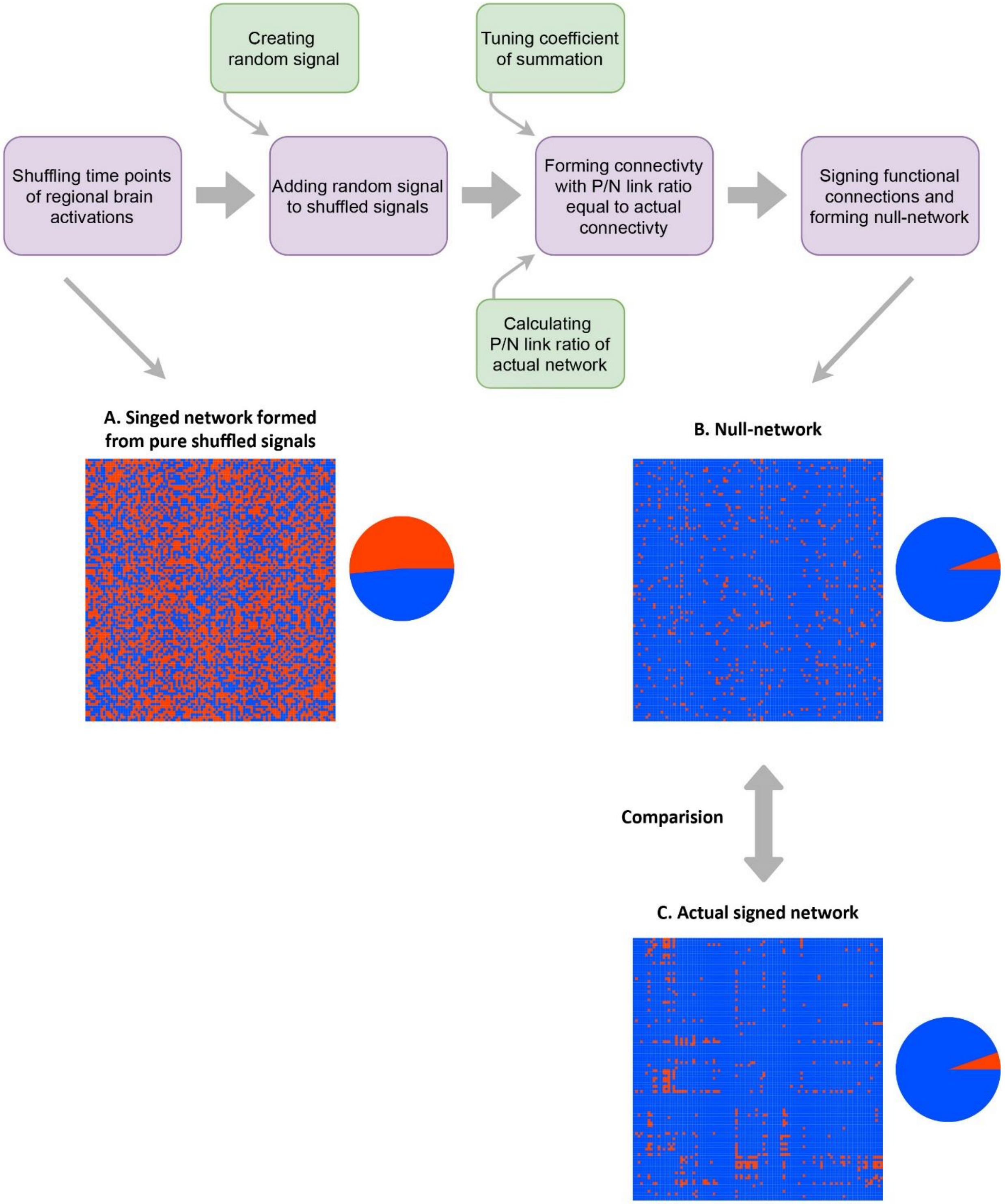


**Figure S7: Illustration of null-network formation for a subject.** Top: procedure of null-network formation. Below: connectivity matrices and signed links percentages for a subject before and after adjusting process. Image plots show signed connectivity matrices and piecharts demonstrate percentages of positive and negative links. Blue and red colors indicate positive and negative links, respectively.

### Supplementary tables:

**Table S1**: Brain regions in which the distribution of negative degrees significantly differ from the whole brain distribution. Columns of the table describe cortical and statistical characteristics of the distributions. Highlighted rows indicate significant ROIs with FDR p-values lower than 0.05.

| | **ROI** | **Canonical Network** | **Hemisphere** | **Rate Parameter** | **p-value** | **FDR p-value** |
|---|---|---|---|---|---|---|
| 1 | Par_2 | Default Mode | Left | 0.083 | 0.013 | 0.111 |
| 2 | pCunPCC_1 | Default Mode | Left | 0.078 | 0.013 | 0.111 |
| 3 | pCunPCC_2 | Default Mode | Left | 0.074 | 0.002 | 0.050 |
| 4 | PFC_3 | Default Mode | Left | 0.080 | 0.048 | 0.221 |
| 5 | PFC_5 | Default Mode | Left | 0.082 | 0.015 | 0.112 |
| 6 | Post_1 | Dorsal Attention | Left | 0.238 | 0.038 | 0.214 |
| 7 | Post_2 | Dorsal Attention | Left | 0.119 | 0.022 | 0.143 |
| 8 | FrOperIns_2 | Ventral Attention | Left | 0.077 | 7.00E-05 | 0.002 |
| 9 | Med_1 | Ventral Attention | Left | 0.491 | 5.00E-05 | 0.002 |
| 10 | ParOper_1 | Ventral Attention | Left | 0.112 | 0.048 | 0.221 |
| 11 | SomMot_6 | Somatomotor | Left | 0.313 | 0.044 | 0.221 |
| 12 | Cont_Par_1 | Frontoparietal | Right | 0.089 | 1.00E-05 | 0.001 |
| 13 | Cont_Par_2 | Frontoparietal | Right | 0.090 | 0.015 | 0.112 |
| 14 | Cont_PFCmp_1 | Default Mode | Right | 0.293 | 0.006 | 0.076 |
| 15 | Par_1 | Default Mode | Right | 0.087 | 0.006 | 0.076 |
| 16 | pCunPCC_2 | Default Mode | Right | 0.065 | 0.001 | 0.023 |
| 17 | PFCdPFCm_1 | Default Mode | Right | 0.086 | 0.044 | 0.221 |
| 18 | Temp_3 | Dorsal Attention | Right | 0.276 | 0.003 | 0.058 |
| 19 | Post_1 | Dorsal Attention | Right | 0.335 | 0.031 | 0.184 |
| 20 | FrOperIns_1 | Ventral Attention | Right | 0.092 | 0.008 | 0.086 |
| 21 | TempOccPar_2 | Ventral Attention | Right | 0.094 | 0.004 | 0.062 |
| 22 | 7Networks_RH_SomMot_8 | Visual | Right | 0.306 | 0.020 | 0.133 |

**Table S2:** Study separated demography and imaging protocols.

| Study | Dataset | Final selected subjects | Age(mean ± SD) | FIQ(mean ± SD) | VIQ(mean ± SD) | PIQ(mean ± SD) | Voxel size(mm) | Anatomical Flip Angle(Deg) | Functional Flip Angle(Deg) | Anatomical Echo Time(ms) | Functional Echo Time(ms) | Anatomical Repetition Time(ms) | Functional Repetition Time(ms) |
|---|---|---|---|---|---|---|---|---|---|---|---|---|---|
| ETH Zürich | ABIDE II | 13 | 25 ± 3 | 118 ± 9 | 114 ± 13 | 113 ± 10 | 3.0×3.0×3.0 | 8 | 90 | Shortest | 25 | 8.4 | 2000 |
| University of Utah School of Medicine | ABIDE I | 20 | 23 ± 4 | 113 ± 13 | 112 ± 13 | 117 ± 13 | 3.4×3.4×3.0 | 9 | 90 | 2.91 | 28 | 2300 | 2000 |
| University of Michigan | ABIDE I | 5 | 22 ± 5 | 115 ± 4 | 117 ± 7 | 112 ± 5 | 3.348×3.348×3.0 | 90 | 90 | 5.7 | 30 | 250 | 2000 |
| NYU Langone Medical Center | ABIDE I | 19 | 24 ± 4 | 117 ± 10 | 117 ± 12 | 114 ± 9 | 3.0×3.0×3.0 | 7 | 90 | 3.25 | 15 | 2530 | 2000 |
| Total | | 57 | 24 ± 4 | 116 ± 11 | 114 ± 13 | 113 ± 10 | | | | | | | |
